# The genetic architecture of human programmed stop codon readthrough

**DOI:** 10.64898/2026.07.14.738498

**Authors:** Ignasi Toledano, Fran Supek, Ben Lehner

## Abstract

Programmed translational readthrough produces C-terminally extended protein isoforms via decoding of stop codons by near-cognate tRNAs. Human genes experimentally validated as readthrough targets share a CUAG motif downstream of a UGA stop codon. However, the full sequence determinants of readthrough efficiency, how they combine, and how generalisable they are across genes remain largely unexplored. Here we use deep mutational scanning to quantify ∼1,400 sequence variants for each of the three examples of human readthrough in the genes *AQP4*, *MAPK10* and *OPRK1*. In addition to the core CUAG motif, mutations that modulate readthrough elements extend up to +27 nucleotides downstream of the stop codon and across six codons (18 nucleotides) upstream. For the downstream sequence, an additive model with a sigmoidal global epistasis function captures most of the within-gene readthrough variance for double mutants (R²=0.84-0.96), with additional contributions from a small number of strong pairwise interactions. Mutational effects nonetheless generalise poorly between genes: only the immediate -3 to +4 nucleotide window shows consistent behaviour, while mutations in more distal positions have context-dependent effects. Combinatorial assembly of sequence blocks from different genes into chimeras reveals strong interactions (epistasis) between sequences upstream and downstream of the stop codon. This study provides comprehensive quantitative maps of the sequence determinants of human programmed readthrough and suggests that three examples of programmed readthrough are located on distinct local fitness peaks, each defined by different upstream and downstream architectures built around a shared CUAG core motif.

## Introduction

When a stop codon enters the ribosomal A site, the translation termination machinery assembles to halt elongation, release the nascent polypeptide, and initiate ribosome recycling. Stop codons are recognized by the release factor eRF1, which is delivered to the ribosome in complex with the GTP-bound release factor eRF3<u>(Alkalaeva et al. 2006)</u>. Stop codon recognition by eRF1 induces conformational rearrangements within the ribosome that trigger eRF3-mediated GTP hydrolysis, leading to peptide release<u>(Alkalaeva et al. 2006;</u> <u>Brown et al. 2015; Lawson et al. 2021)</u>. However, translation termination is not 100% efficient and stop codons can be decoded by near-cognate tRNAs, which insert an amino acid and allow elongation to continue until the next in-frame termination signal. This phenomenon is referred to as translational stop codon readthrough<u>(Dabrowski et al. 2015;</u> <u>Palma and Lejeune 2021)</u>.

Basal readthrough occurs spontaneously at extremely low frequencies (typically <0.1%)<u>(Floquet et al. 2012; Palma and Lejeune 2021; Toledano et al. 2024)</u>. Nevertheless, some genes have specific sequence contexts around the stop codon that reduce termination fidelity and promote readthrough, resulting in the synthesis of substantial levels of C-terminally extended protein isoforms<u>(Pelham 1978; Brown et al. 1996; Firth et al. 2011;</u> <u>Robinson and Cooley 1997; Lin et al. 2007; Jungreis et al. 2011; Dunn et al. 2013;</u> <u>Prieto-Godino et al. 2016; Loughran et al. 2014; Eswarappa et al. 2014; Schueren et al.</u> <u>2014; Loughran et al. 2018)</u>. These events are known as programmed readthrough, and it has been documented across diverse evolutionary lineages.

Programmed readthrough was first described in viruses, where it contributes to the optimization of genome coding capacity<u>(Weiner and Weber 1971; Pelham 1978; Dreher and</u> <u>Miller 2006; Firth and Brierley 2012)</u>. Compact viral genomes must fit within the constraints of small capsids while still encoding the full repertoire of proteins required for the viral life cycle. In murine leukemia gammaretrovirus (MuLV), readthrough enables the expression of two proteins from a single mRNA, translated as an extended polyprotein that is subsequently proteolytically cleaved<u>(Philipson et al. 1978; Yoshinaka et al. 1985)</u>. Similarly, in several plant viruses, readthrough is used to express extended isoforms of their replicase proteins<u>(Pelham 1979; Dreher and Miller 2006)</u>. Beyond viruses, extensive readthrough has been reported in some *Drosophila* species, where comparative genomics and ribosome profiling<u>(Ingolia et al. 2009)</u> studies predicted around 300 candidate readthrough genes, with tissue-specific readthrough for some of them<u>(Robinson and Cooley 1997; Lin et al. 2007;</u> <u>Jungreis et al. 2011; Dunn et al. 2013; Chen et al. 2020; Makarova et al. 2026)</u>. One striking example is gene *Dsec*Ir75a, which undergoes ∼90% readthrough exclusively in neurons<u>(Prieto-Godino et al. 2016)</u>. Finally, a combination of comparative genomics and motif-based analyses suggested the existence of readthrough in human genes<u>(Jungreis et</u> <u>al. 2011; Lindblad-Toh et al. 2011; Loughran et al. 2018)</u>, and subsequent experimental work validated several cases<u>(Yamaguchi et al. 2012; Schueren et al. 2014; Loughran et al. 2014;</u> <u>Eswarappa et al. 2014; Stiebler et al. 2014; Loughran et al. 2017; Loughran et al. 2018;</u> <u>Yordanova et al. 2018; Singh et al. 2019; Manjunath et al. 2020)</u>. Excluding the special case of selenocysteine incorporation<u>(Gonzalez-Flores et al. 2013)</u>, 14 human genes with programmed readthrough have been experimentally validated, with diverse functional consequences of the extended isoforms. These include the acquisition of organelle-targeting signals<u>(Schueren et al. 2014; Stiebler et al. 2014)</u>, altered subcellular localization near the vasculature<u>(De Bellis et al. 2017; Mohaupt et al. 2023)</u>, protein binding inhibition<u>(Singh et al.</u> <u>2019)</u>, regulation of translation<u>(Yordanova et al. 2018)</u> and contributions to maintaining mitochondrial membrane potential<u>(Manjunath et al. 2020)</u>. In some cases, however, the functional consequences - if any - remain unknown<u>(Loughran et al. 2014)</u>.

The sequence encoding the readthrough signal also varies across genes. First, there is a group of genes where the sequence determinants of readthrough are unknown (e.g., *HBB*, *AMD1*). In a second group, readthrough is promoted by *trans*-acting factors binding specific sequence elements downstream of the stop codon (e.g., *VEGFA*, *AGO1*). For a third group—including *AQP4*, *MAPK10*, *OPRK1*, *OPRL1*, *VDR*, *MDH1*, and *LDHB* - readthrough is promoted by a shared CUAG motif located downstream of a UGA stop codon. Notably, despite sharing this motif, these genes exhibit distinct readthrough efficiencies, indicating that the readthrough motif is only partially mapped and that additional nucleotides (nts) must modulate it. A recent study tested a limited number of mutations in *MDH1*, *LDHB*, and *AQP4*, identifying some mutations beyond the CUAG motif with significant readthrough modulatory effects<u>(Smoljanow et al. 2025)</u>. However, the limited throughput of the approach (∼25 variants tested per gene) precluded a comprehensive and quantitative understanding of the sequence determinants governing readthrough.

In addition to programmed readthrough, readthrough of stop codons can also be promoted by small molecule drugs<u>(Dabrowski et al. 2018; Li et al. 2023)</u>. We recently used pooled selection and sequencing experiments using libraries of thousands of different premature stop codons to explore how sequence context modulates readthrough promoted by small molecule drugs. In addition to stop codon identity, we identified strong effects of the 3nts upstream and the 3nts downstream of the stop codon, allowing interpretable models to be trained that predict the quantitative readthrough of stop codons across the human genome by each drug<u>(Toledano et al. 2024)</u>.

However, for detectable readthrough in the absence of drugs, stronger sequence contexts are required and these contexts have not been extensively studied using mutagenesis. As such, our understanding of how programmed readthrough is encoded in sequence is incomplete and basic questions such as which positions are involved, how residues in different positions combine and how generalizable mutation effects are across genes, are still largely unaddressed.

Here, we use mutagenesis to systematically dissect the genetic architecture of three examples of programmed translational readthrough in human genes. For each of three examples (*AQP4*, *MAPK10*, and *OPRK1*) we quantified the effect on readthrough of >1,350 diverse mutations. To our knowledge, this constitutes the first comprehensive deep mutational scanning dataset for programmed readthrough. We identified individual mutations with strong effects distributed broadly across the sequence, extending up to 27 nucleotides away from the stop codon. Our double-mutant strategy further allowed us to assess how mutations combine and to fit gene-specific predictive models of readthrough. Finally, we explored how mutations generalise across genes and found that programmed readthrough is shaped by gene-specific architectures built around a shared CUAG core motif.

## Results

### Deep mutagenesis of programmed readthrough

We designed libraries mutating the sequence around the stop codons of four examples of human stop codon readthrough validated in multiple studies - *AQP4, MAPK10, OPRK1* and *OPRL1*. PhyloP and phastCons scores calculated across 100 vertebrate species show that the readthrough extension has nucleotide conservation comparable to that of the main open reading frame (ORF) **(Fig. 1a, Extended Data Fig. 1a)**, whereas conservation drops sharply downstream of the second stop codon (3’ UTR), suggesting functional relevance of the readthrough extension. In addition, these three extensions have previously been reported to have PhyloCSF scores typical of protein coding regions<u>(Loughran et al. 2014)</u>.

**Figure 1.**
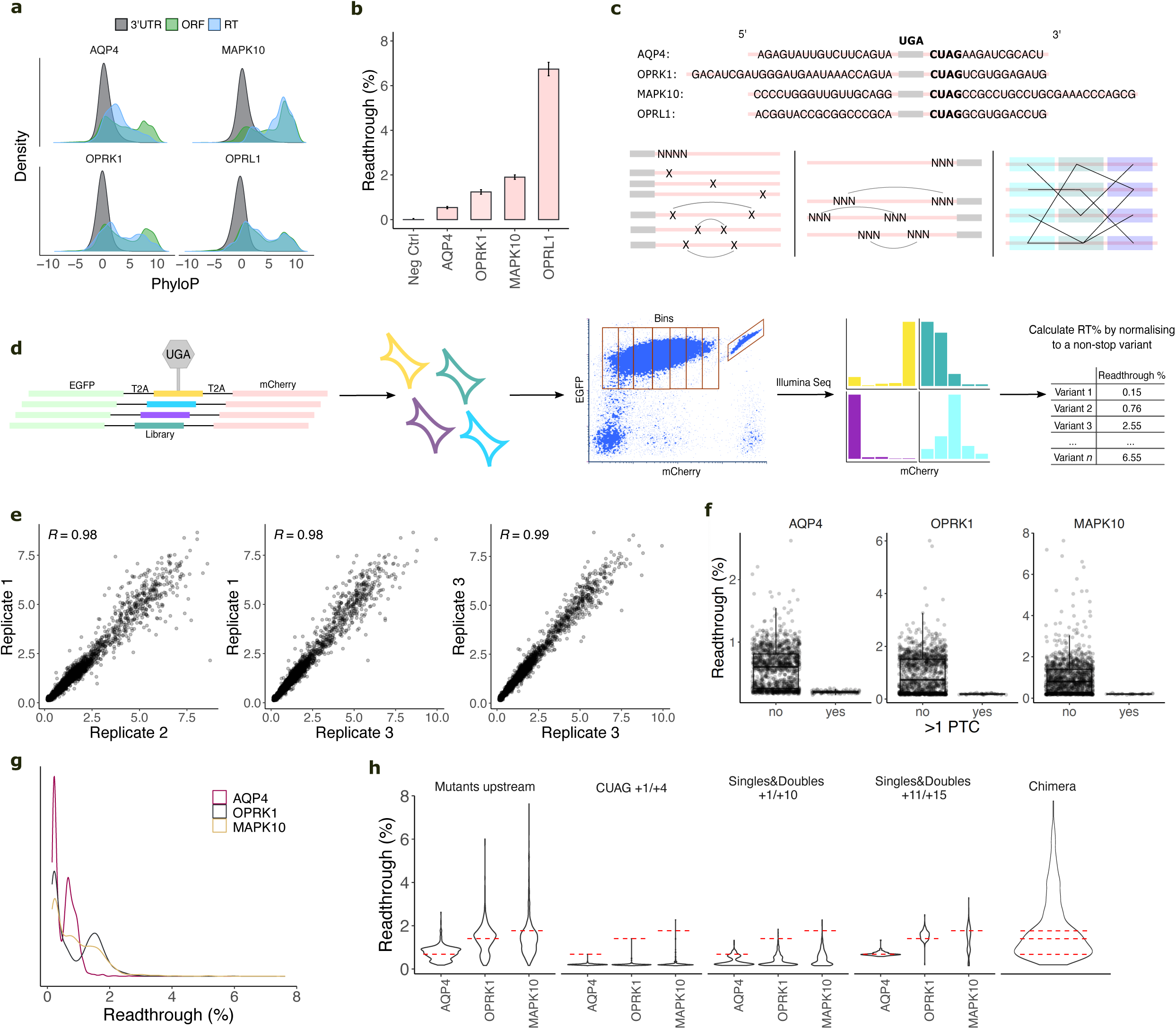
Deep mutagenesis of four human programmed readthrough genes. **a,** Distribution of PhyloP scores across the open reading frame (ORF), readthrough extension (RT) and the rest of the 3’UTR across the *AQP4*, *MAPK10*, *OPRK1* and *OPRL1* genes. **b,** Readthrough of the four genes’ *WT* relative to a non-stop variant. A stop codon variant eliciting strong translation termination was used as negative control. Error bars indicate standard deviation across 3 replicates. **c,** Mutagenesis strategy. The *WT* sequences of the four genes (with the shared UGACUAG motif highlighted in bold) and the three mutagenesis strategies are shown. Downstream, we tested all single and double mutants (*left*). Upstream, we designed all single codon mutants in the position right upstream of the stop codon plus sampled ∼500 double or triple mutants per gene, across all upstream positions (*center*). Finally, we designed chimeric variants where each gene was divided into six chunks and these were randomly combined preserving the position order (*right*). **d,** Experimental design, 5980 mutations around the *AQP4*, *MAPK10, OPRK1* and *OPRL1* stop codon were cloned in the readthrough fluorescent reporter<u>(Toledano et al. 2024)</u> and integrated into the genome of a HEK293T landing pad human cell line<u>(Matreyek et al. 2020)</u> **(Methods)**. Cells were sorted in eight bins based on mCherry fluorescence, each bin was Illumina-sequenced and readthrough percentages were calculated from the distribution of reads across bins normalized to a non-stop variant. **e,** Correlations of readthrough scores for all variants (with >100 reads) across three replicates. **f,** All negative control variants with two stop codons display ∼0% readthrough, confirming that the mCherry fluorescent signal arises exclusively from readthrough. The top and bottom sides of the box are the lower and upper quartiles. The box covers the interquartile interval, where 50% of the data are found. The horizontal line that splits the box in two is the median. **g,** Readthrough distribution across genes for all variants (with >100 reads). **h,** Readthrough distribution across the five different mutation classes. The dashed red lines indicate the readthrough of each gene’s *WT*.

*AQP4*, *MAPK10*, *OPRK1* and *OPRL1* have the same CUAG tetranucleotide downstream of the UGA stop codon. However they were previously reported to have different readthrough efficiencies<u>(Loughran et al. 2014)</u>. Using serial truncations, the authors defined the positions around the stop sufficient for wild-type (*WT*)-levels of readthrough, which ranged between 18-27 nts upstream and and 15-27 nts downstream (18, 18, 27 and 18 nts upstream, and 15, 27, 15 and 15 nts downstream for *AQP4*, *MAPK10*, *OPRK1* and *OPRL1*, respectively) (**Fig. 1c)**. We adopted these sequence boundaries throughout this study.

We first quantified the readthrough of these sequences in our experimental platform<u>(Toledano et al. 2024)</u> **(Fig. 1b, Methods)**. Each variant was cloned into a dual fluorescent protein reporter, where an upstream enhanced green fluorescent protein (EGFP) controls for variable expression and readthrough causes translation of a downstream mCherry protein, and performed single-copy genomic integration into a HEK293T landing pad cell line<u>(Matreyek et al. 2020)</u>. Readthrough was calculated by normalizing the mCherry/EGFP ratio of each variant to a non-stop control variant representing 100% efficient translation. Readthrough scores were 0.5%, 1.2%, 1.9% and 6.4% for *AQP4*, *MAPK10, OPRK1* and *OPRL1*; respectively. A stop codon eliciting strong translation termination (*TP53* R213X) was used as negative control<u>(Toledano et al. 2024)</u>.

After validating that sequences beyond the CUAG motif further influence readthrough, we designed mutagenesis libraries to explore their contribution **(Fig. 1c)**. Upstream of the UGA stop codon, we designed all individual codon variants in the -1 codon (61 sequences per gene), along with a sparse subsampling of two and three codon variants across the six codons upstream (∼500 sequences per gene). Downstream, we fully randomized the CUAG tetranucleotide (4^4^ = 256 sequences), tested all single mutations in between nucleotide positions +1 and +15 (45 sequences), and tested all double mutations in two groups (positions 1 to 10 and 11 to 15; 495 sequences). Finally, we divided each gene into six sequence blocks (three upstream and three downstream, 5-9 nts each depending on the *WT* sequence length) and combined them across genes while preserving block order (ie. block 1 always tested in position 1). Specifically, we evaluated all possible downstream block combinations in combination with the *WT* upstream sequences and, conversely, all possible upstream block combinations against the *WT* downstream sequences, for a total of 448 chimeric variants. As a control, we included a non-stop variant representing 100% efficient translation to which all variants in the assay were normalized **(Methods)**.

We cloned the library into our readthrough reporter<u>(Toledano et al. 2024)</u> and stably integrated it into the genome of the HEK293T landing pad cells. We combined fluorescence sorting and Illumina sequencing to quantify readthrough efficiencies for each variant, expressed as the percentage of expression relative to the non-stop control variant **(Supplementary Table 1, Methods)**. Briefly, readthrough scores were calculated by sorting cells into eight bins according to mCherry fluorescence, sequencing each bin independently, and determining the distribution of reads across bins **(Fig. 1d, Extended Data Fig. 1b, Methods)**.

Readthrough scores were highly correlated across the three replicates (*R* = 0.98-0.99) **(Fig. 1e)**. Of the 5980 variants designed, we retrieved 5036 with ≥100 reads (*n*=1367, 1394, 1370 and 482 for *AQP4*, *MAPK10*, *OPRK1* and *OPRL1*; the remaining 423 sequences are the chimeric variants). Most *OPRL1* variants were lost during library integration, and we therefore excluded this gene from all downstream analyses except for the chimera variants, where *OPRL1* variants were recovered at a frequency comparable to the other three genes. As an internal control, we used 120 variants where the mutation introduces a second stop codon which should reduce the readthrough efficiency to close to 0%. Across all three genes we observed the expected behaviour for these variants **(Fig. 1f)**, supporting readthrough as the main driver of signal in our assay.

Throughout the paper, positions upstream of the stop codon are given in codon units, starting at -1 (the codon immediately upstream), whereas positions downstream are given in nucleotide units, starting at +1 (the nucleotide immediately downstream). This dual nomenclature reflects the mutagenesis design: upstream mutations were introduced at the codon level, downstream mutations at the nucleotide level.

### Distributions of readthrough mutational effects across genes

The distribution of readthrough scores differs across genes **(Fig. 1g)**. *AQP4* scores range from 0-2.62% with the 75th percentile at 0.77%. In contrast, *OPRK1* and *MAPK10* displayed distributions shifted to higher values, with 0-6% and 0-7.6% ranges, and 75th percentiles at 1.48% and 1.40%, respectively. Interestingly, the number of mutations that increase readthrough relative to the *WT* inversely relates to the *WT* readthrough level **(Fig. 1h)**. Specifically, *AQP4*, *OPRK1*, and *MAPK10* exhibit *WT* readthrough rates of 0.68%, 1.39%, and 1.76%, respectively, whereas the number of mutations that enhance readthrough are 247, 182, and 77, respectively (FDR = 0.05, two-sided T-test, compared to *WT*).

Classification of variants by mutation type revealed three main findings **(Fig. 1h)**. First, variants in all mutated regions can alter readthrough (FDR = 0.05). Second, mutations in different regions had markedly different effect sizes. Specifically, upstream and chimeric mutants registered the largest readthrough-increasing effects. Third, the direction of mutational effects differs in the different regions. In particular, mutations in the four nucleotides immediately downstream of the stop codon consistently decrease readthrough below *WT* levels, as opposed to other regions where mutations also enhance readthrough. In the following sections we further examine each mutation class.

### Influence of the downstream sequence

We first focused on the readthrough-modulatory potential of the four nucleotides downstream of the stop codon, where the *WT* sequence is CUAG in all three genes and for which we tested all 4^4^=256 possible combinatorial variants. CUAG has been reported as a strong readthrough stimulator<u>(Loughran et al. 2014; Loughran et al. 2018)</u>, consistent with the presence of this motif in all four genes. In line with this, 764 of the 765 mutations at positions +1 to +4 reduced readthrough relative to *WT*, with only CUAA increasing *MAPK10* readthrough from 1.76% to 2.28% (FDR=4e-04) **(Fig. 2a)**. We observed a Hamming distance (HD)-dependent effect, whereby the greater the number of mutations from the *WT* sequence, the larger the decrease in readthrough up to HD=3 *vs* HD=4 variants, for which readthrough is ∼0 **(Fig. 2a**, **Extended Data Fig. 1d)**. Variants with more than one mutation mostly abolished readthrough across all genes, whereas a small number of double mutants retained low levels in *MAPK10*. Across all three genes, a subset of single mutants also retained some readthrough, and these showed high correlation across genes (R = 0.69, R =0.74, R=0.74; for *AQP4-OPRK1*, *AQP4-MAPK10* and *OPRK1-MAPK10*, **Extended Data Fig. 1d**). A common feature among all readthrough variants in the dataset was the presence of the CU dinucleotide in positions +1 and +2, highlighting its essential role in enabling readthrough in these sequence contexts **(Fig. 2a,b)**.

**Figure 2.**
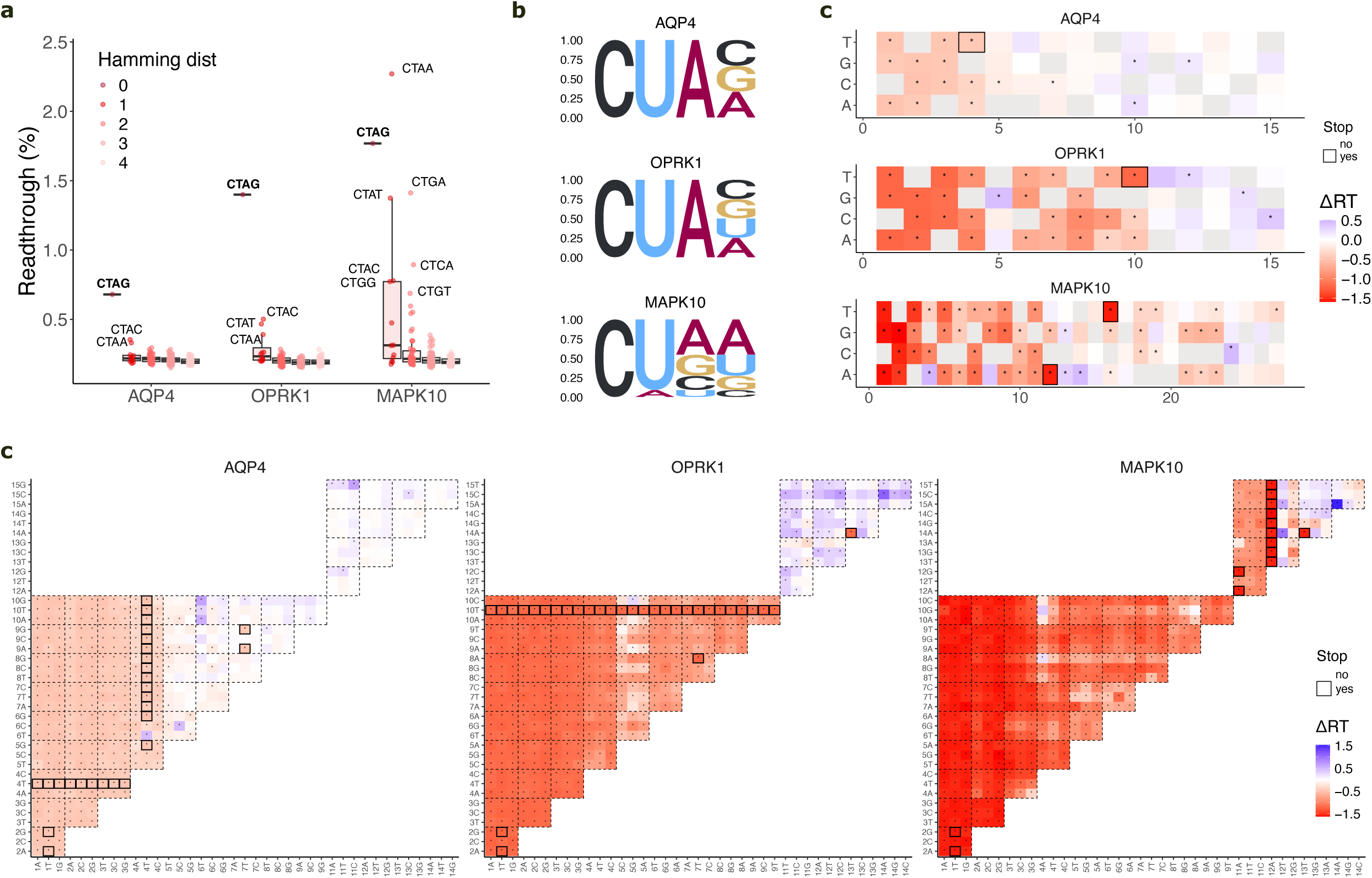
Readthrough modulation by the downstream sequence. **a,** All mutants in the complete mutagenesis of +1 to +4 positions (CUAG) are shown. Color legend indicates the hamming distance from the CUAG *WT* (bold). The sequence of the highest readthrough variants in each gene is shown. **b,** Sequence logos for the CUAG-randomized mutants with the highest readthrough per gene (RT>0.3%, n=3; RT>0.3%, n=3; RT>0.5%, n=11; for *AQP4*, *OPRK1* and *MAPK10*; respectively). **c,** Delta readthrough (ΔRT, relative to *WT*) of all single mutants downstream of the stop codon, from +1 to +15 (*AQP4*, *MAPK10*) or to +27 (*MAPK10*). Grey indicates the *WT* nucleotide. Asterisks indicate that delta readthrough is significantly different from zero (FDR < 0.05, two-sided T-test). The black square indicates that the mutation introduced a second stop codon in frame, and are therefore considered negative controls. **d,** Same as **c** but for all double mutants downstream of the stop codon.

Beyond the fully randomized CUAG context, the library included all single mutants spanning +1 to +15 (*AQP4*, *OPRK1*) or +1 to +27 (*MAPK10*), and all double mutants spanning +1 to +15 across all genes. For each mutant we calculated the delta readthrough (ΔRT) relative to *WT* and observed that most mutations in this region reduced readthrough. At FDR=0.05, 54%, 80%, and 88% of variants decreased readthrough in *AQP4*, *OPRK1*, and *MAPK10*, respectively **(Fig. 2c,d)**. In contrast, a smaller fraction increased readthrough, accounting for 3%, 6%, and 2% of mutations in the same genes.

The readthrough signal extends over a longer sequence than previously appreciated, with distal mutations (+5 to +15) modulating readthrough across all three genes (relative to *WT*; FDR = 0.05) **(Fig. 2c)**. In *MAPK10*, where a longer region was interrogated, these effects extend up to position +27. Notably, distal mutations can exhibit large effect sizes, in some cases comparable to CUAG variants, such as mutations in nucleotides +5 to +10 in *OPRK1* and in +5 to +12 in *MAPK10* (up to 1.2% decrease in both genes). As an illustrative example, the mean ΔRT across the six +8 and +9 mutants is -0.75% and -0.82% for *OPRK1* and *MAPK10*; respectively. Furthermore, 6 of 9 *MAPK10* mutations at positions +21 to +23 reduce readthrough by more than 0.5%, highlighting the contribution of long-range effects.

Another important observation is that the effect of mutations is strongly gene-dependent. For example, at position +5 the nucleotide with highest readthrough is A, G and C for *AQP4*, *OPRK1*, and *MAPK10*, respectively. Similarly, at position +8 the highest readthrough nucleotide is T for *OPRK1* but A for *MAPK10*, whereas for *AQP4* there are no significant differences across the four variants (one-way ANOVA test) **(Fig. 2c)**. In line with this, in 6 of 8 positions within +5 to +12, the nucleotide yielding the highest readthrough differs between *OPRK1* and *MAPK10* **(Fig. 2c)**.

This sequence context specificity is further reflected in the number of positions that modulate readthrough and in the direction of their effects. In *AQP4*, only 7 of the 33 single mutants within the +5 to +15 region significantly modulate readthrough, of which 3 increase it (FDR=0.05). In contrast, 18 and 24 of the +5 to +15 mutants significantly modulate readthrough in *OPRK1* and *MAPK10*, respectively, although a smaller fraction increase it (4 of 18 and 3 of 24, all FDR=0.05). This, along with several other significant effects in positions +15 to +27 in *MAPK10*, shows that the higher *WT* readthrough levels in *OPRK1* and *MAPK10* compared to *AQP4* depend on readthrough-promoting sequences throughout a longer downstream sequence.

Finally, the double mutants provide additional insight (further analysed in the next section, ‘Interpretable models to predict readthrough’) **(Fig. 2d)**. An interesting case is the *AQP4* 4G>T_6A>T mutation, which is the only *AQP4* variant carrying a CUAG-disrupting mutation that increases readthrough above WT (ΔRT = 0.5%, FDR = 3e-03). Beyond this case, *AQP4* contains 8 additional readthrough-increasing double mutants involving position +10, consistent with the readthrough-promoting effect of +10 mutations seen at the single-mutant level. In contrast, *OPRK1* and *MAPK10* each contain only one readthrough-increasing double mutant within positions +1 to +10 (5T>A_10G>C and 4G>A_8C>A, respectively). In the +11 to +15 region, however, all three genes contain several readthrough-increasing double mutants, with *OPRK1* showing the largest number (27, compared with 6 in *AQP4* and 8 in *MAPK10*). The strongest examples per gene are 11G>C_15T>G in *AQP4* (ΔRT = 0.65%), 14T>A_15G>C in *OPRK1* (ΔRT = 1.1%), and 14T>A_15G>A in *MAPK10* (ΔRT = 1.5%).

Overall, a consistent pattern is seen across the three genes: the frequency of readthrough-enhancing mutations increases from proximal to distal positions **(Fig. 2c,d)**. Although the immediate downstream sequences are optimal for readthrough in all three genes, multiple distal mutations can further increase it.

### Interpretable models to predict readthrough

We next asked to what extent simple, interpretable additive models could predict readthrough based on the downstream sequence. For each gene, we fitted a model that predicts the readthrough of a variant from its set of mutations relative to a reference sequence. The model has two components: an additive genetic-trait term, in which each nucleotide at each position contributes a fixed coefficient, and a sigmoidal global epistasis function<u>(Domingo et al. 2019)</u> that maps this latent additive trait onto the measured readthrough<u>(Toledano et al. 2024)</u>, capturing the non-linearity arising from the readout’s lower bound at 0% **(Methods, Extended Data Fig. 1e)**.

We trained the model on the downstream single and double mutants of each gene. Each single mutation was observed in an average of 27 (for positions 1 to 10) and 12 (for positions 11 to 15) different sequences (backgrounds), providing robust background-averaged estimates of individual mutational effects. To ensure a more-balanced representation of training and test data across the sequence, we excluded the higher-order (third- and fourth-order) CUAG mutants. The *AQP4 WT* was used as a common arbitrary reference sequence in all three models, so that the resulting coefficients are directly comparable between genes. Performance was evaluated by 10 rounds of cross-validation with a 90-10 train-test split. The held-out predictions correlated strongly with the measured readthrough values, yielding R² = 0.89, 0.96 and 0.84 for *AQP4*, *OPRK1* and *MAPK10*, respectively, showing that a simple combination of additivity and a non-linear global epistasis function captures the majority of the readthrough variance for each gene **(Fig. 3a, Extended Data Fig. 1e)**.

**Figure 3.**
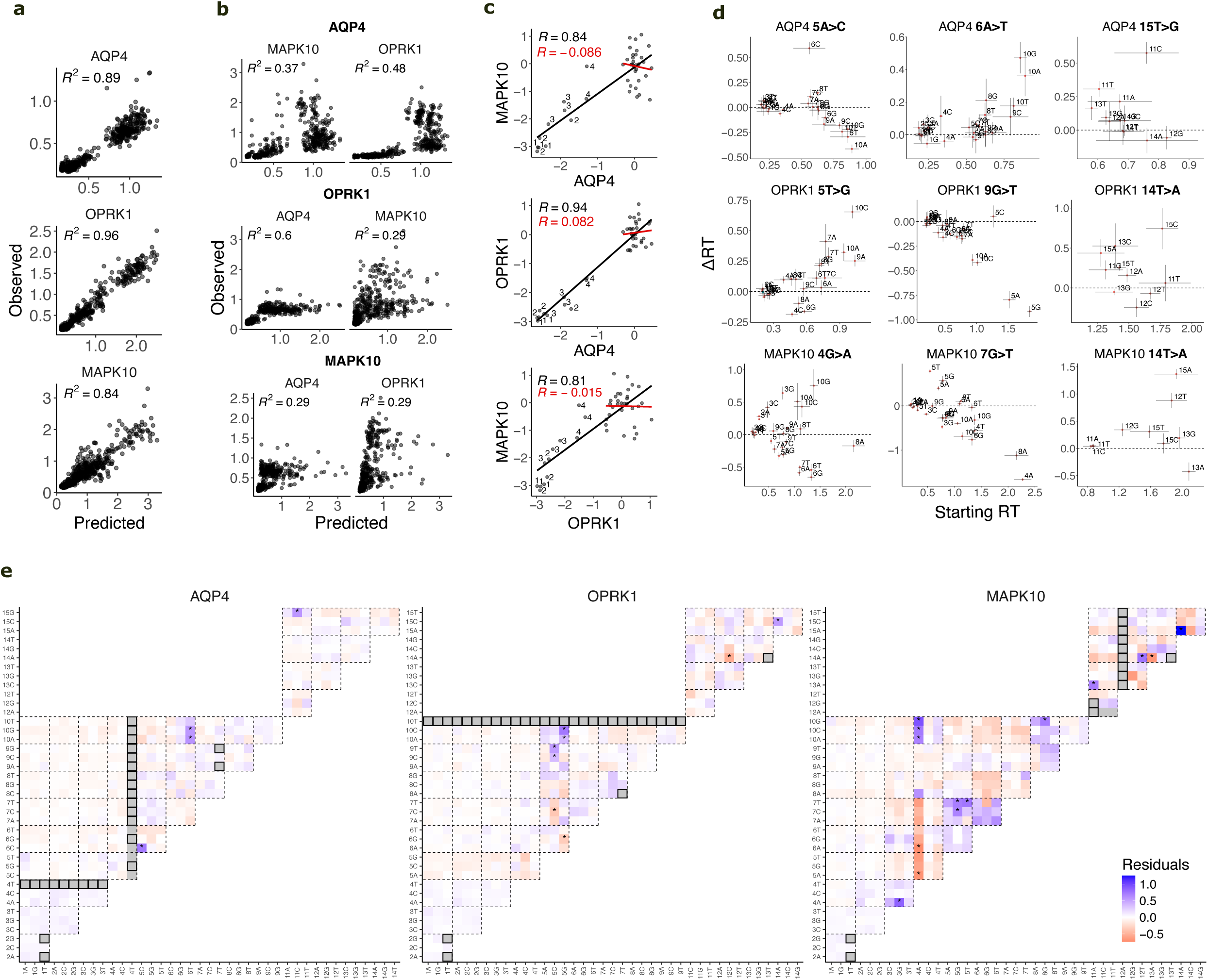
Sequence additivity and epistasis in the downstream sequence. **a,** Predicted *vs* observed readthrough scores for gene-specific models that use the sequence downstream of the stop codon to predict the readthrough efficiency of a variant. Cross-validated *R^2^* is shown. **b,** Each gene-specific model (in bold) was tested on the two other genes. Cross-validated *R^2^* is shown. **c,** Correlation of the models’ coefficients across genes. Coefficients belonging to mutations in positions +1 to +4 have the position annotated. *R^2^*s considering all positions (black) or excluding the +1 to +4 positions (red) are shown. **d,** For a given individual mutation (indicated in the title), the readthrough of each background (X axis, Starting RT) and the delta readthrough upon introducing the mutation (Y axis, ΔRT) are shown. The identity of the background mutations are indicated. Error bars indicate the standard deviation across the three replicates of the single mutant background (horizontal) and the double mutant (vertical). The dashed line defines the boundary between negative and positive effects. **e,** Residuals (predicted - observed) for all single and double mutants downstream. Asterisks highlight normalized residues greater than three standard deviations of the residuals distribution. Mutations excluded from model fitting are shown in grey, and fall into two classes: a) variants that themselves introduce a second stop codon are additionally outlined in black and b) variants in which one of the constituent single mutations would create a stop codon in most other backgrounds (but does not do so in this specific background) are shown in grey only, without the black outline.

We next evaluated each model’s ability to predict readthrough in the other two genes, asking to what extent mutational effects are preserved across sequence contexts. Performance dropped substantially across all three models. The *AQP4*-trained model achieved R² = 0.37 on *MAPK10* and 0.48 on *OPRK1*; the *OPRK1*-trained model achieved R² = 0.6 on *AQP4* and 0.29 on *MAPK10*; and the *MAPK10*-trained model achieved R² = 0.29 on both *AQP4* and *OPRK1* **(Fig. 3b)**. As an additional measure of cross-gene generalisation, we correlated the model coefficients between each pair of genes, obtaining R = 0.84 (*AQP4 vs MAPK10*), 0.94 (*AQP4 vs OPRK1*) and 0.81 (*OPRK1 vs MAPK10*) **(Fig. 3c)**. However, these strong correlations are largely accounted for by the CUAG motif at positions +1 to +4: once these positions are excluded, the correlations drop to R = -0.09 (p = 0.53), R = 0.08 (p = 0.65) and R = -0.02 (p = 0.92), respectively **(Fig. 3c)**. Thus, disruption of the CUAG motif has very similar detrimental effects across genes, whereas mutations at more distal positions have far less conserved effects. Together, these observations are consistent with the discrepancies in mutational effects across genes described above, and quantify the extent to which different sequence backgrounds shape these effects.

### Downstream mutations combine additively with rare epistatic interactions

Even though an additive model with global epistasis explains most of the readthrough variance for each gene, we next leveraged the double mutants to explore potential cases of non-additive interactions between mutations (specific epistasis). Since each double mutant is observed only once, the model can not reliably infer coefficients for pairwise interactions. As an alternative approach, we calculated the *observed vs. predicted* residuals for all double mutants to identify cases with high discrepancy **(Fig. 3e)**. We focused on the double mutants with the largest residuals (defined as those exceeding three standard deviations from the mean of the residual distribution, all significantly different from zero at an FDR < 0.01) to identify candidate cases of specific epistasis. In *AQP4*, the four largest residuals are: 5A>C_6A>C, 6A>T_10C>A, 6A>T_10C>G and 11G>C_15T>G. In *OPRK1*, of the eight largest residuals, six involve a position-5 mutation (5T>C or 5T>G) paired with a mutation at position 6, 7, 9 or 10, while the remaining two involve the distal 14T>A mutation (paired with 12G>C or 15G>C). In *MAPK10*, thirteen strong interactions were identified: six involved the 4G>A mutation, three were pairs between positions 5 and 7 (5C>G_7G>C, 5C>G_7G>T and 5C>T_7G>T), three involve the distal 14T>A mutation (paired with 12C>T, 13C>A or 15G>A) and the last one corresponds to the 11G>A_13C>A pair. Several single mutations recurred across several epistatic interactions, producing residuals of the same sign in some cases and opposite signs in others. For instance, the *OPRK1* mutations 5T>C, 5T>G and 14T>A, and the *MAPK10* mutations 4G>A and 14T>A, produce residuals of both signs depending on the partner mutation. In contrast, the *AQP4* mutation 6A>T and the *MAPK10* mutation 5G>T produce only positive residuals.

To further explore these recurrent mutations, we calculated the change in readthrough (ΔRT) for each mutation across all measured backgrounds (sequences) and plotted it against the background readthrough (starting RT) **(Fig. 3d).** This analysis illustrates how the effects of mutations with specific epistatic interactions vary across backgrounds beyond what global epistasis alone would predict: backgrounds with the same starting readthrough have very different ΔRT upon introducing the mutation. In these cases, neither the magnitude nor the sign of a single mutation’s effect can be inferred without knowing the identity of its partner mutation.

We highlight a few examples of potential interactions, where all reported mutational effects pass FDR < 0.01 in a two-sided T-test comparing the double mutant to the background single mutant across the three replicates **(Fig. 3d)**. One example is the *AQP4* 5A>C mutation, which can have negative, neutral, or positive effects depending on the background. In the 10A background, it reduces readthrough by 0.4%, whereas in the 6C background it increases readthrough by 0.6%. The 15T>G mutation behaves similarly: at the same starting readthrough, it is neutral in the 14A and 12G backgrounds, whereas it increases readthrough by 0.6% when paired with 11C. This is a particularly interesting case given its large distance from the stop codon. In *OPRK1*, the 9G>T mutation switches between neutral and detrimental effects, illustrated by its behaviour across the three position +5 backgrounds, all with very similar starting readthrough. In the 5C background it has no significant effect, whereas in the 5A and 5G backgrounds it reduces readthrough by up to 1%. Finally in *MAPK10*, the 4G>A and 7G>T mutations likewise display all three behaviours depending on the background. For 4G>A, two backgrounds with the same starting readthrough (6G and 10G, both 1.3% RT) show opposite responses: a 0.7% decrease or a 0.8% increase, respectively. Similarly, the 7G>T mutation increases readthrough by 0.8% in the 5T background but decreases it by 1.7% in the 4A background. Interestingly, the 14T>A mutation can either increase readthrough by 1.5% or decrease it by 0.4% depending on the background.

Taken together, our results show that mutations in the downstream sequence mostly combine additively to affect readthrough. However, pairwise epistatic interactions do occur and, as a result of these pairwise interactions, mutational effects substantially alter between one sequence context and another outside of the core CUAG motif. These pairwise interactions - potentially together with higher order interactions that we have not tested - will result in mutations having different effects in different genes. The interacting mutations identified here are adjacent nts and pairs separated by up to 5 nucleotides, suggesting they affect a common molecular mechanism such as a binding site or secondary structure.

### Influence of the upstream sequence

We next explored the sequence upstream of the stop codon, where we generated two groups of mutations. Throughout this section, positions are given in codon units rather than nucleotides, with the codon immediately upstream of the stop defined as position -1. First, we tested all 61 sense codons (along with the three stop codons as controls) at the -1 position. Second, we generated ∼500 higher-order codon mutants per gene: double mutants in *AQP4* and *MAPK10* spanning up to position -6, and triple mutants in *OPRK1* spanning up to position -9.

Mutations upstream of the stop codon drive larger readthrough changes than those downstream **(Fig. 1h)**. Restricting the comparison to double nucleotide mutants (to control for the number of substitutions), the highest readthrough was 1.3%, 2.5% and 3.3% among downstream variants and 2.2%, 5.8% and 7.2% among upstream variants for *AQP4*, *OPRK1* and *MAPK10*, respectively. Mutations in the -1 codon have the strongest effects, although those further upstream still display substantial effect sizes, with some mutations abolishing readthrough completely and others doubling it relative to *WT* **(Fig. 4a)**.

**Figure 4.**
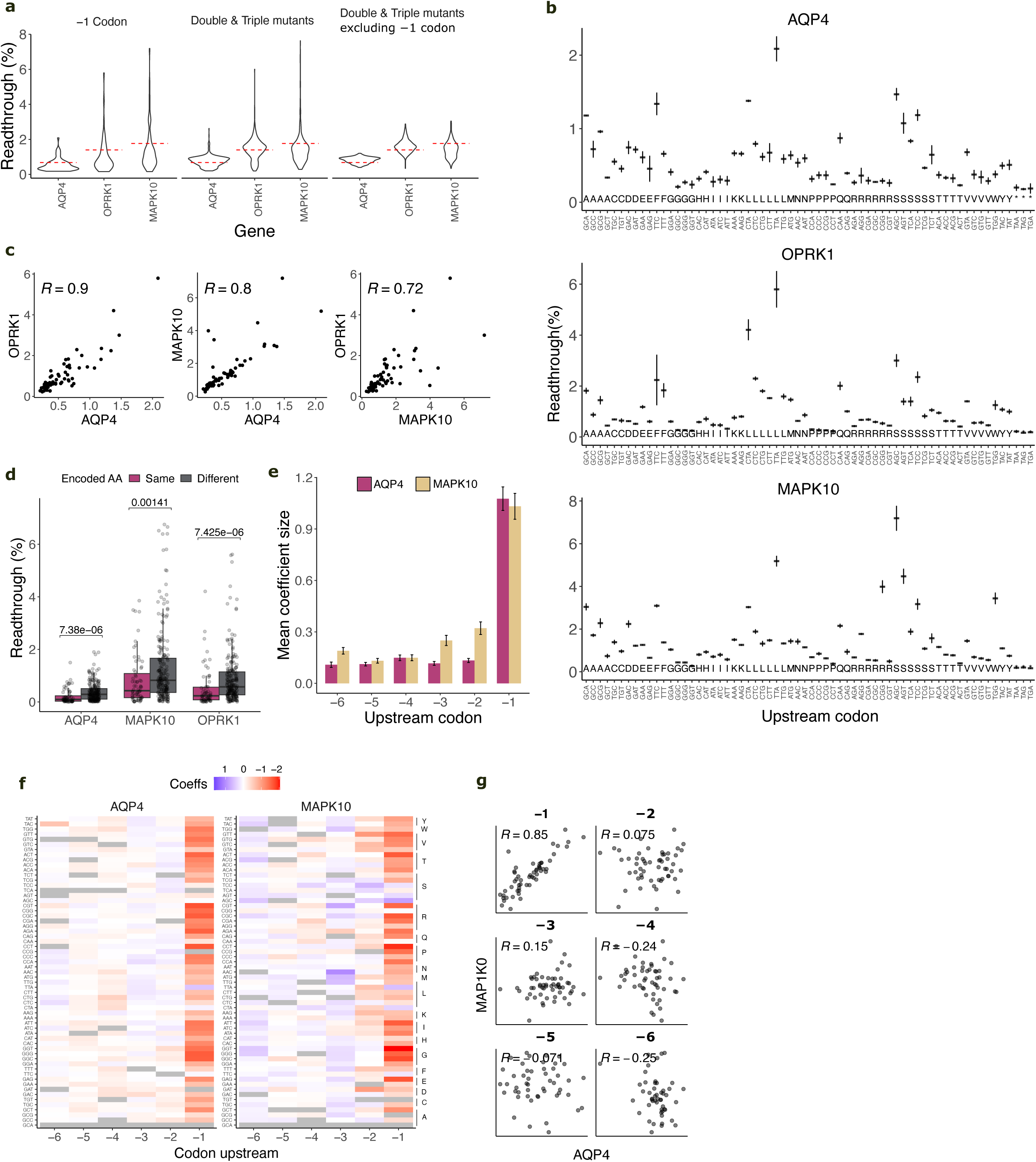
Readthrough modulation by the upstream sequence. **a,** Readthrough distribution across the three different mutation classes of upstream mutants *i)* all -1 codon single mutants, *ii)* double mutants across the -1 to -6 (*AQP4*, *MAPK10*) and triple mutants across the -1 to -9 (*ORPK1*) codons and *iii)* same as *ii* but discarding variants with mutations in the -1 codon. The dashed red lines indicate the readthrough of each gene’s *WT.* **b,** Mean readthrough across the three replicates for each of the -1 codon single mutants in each of the three genes. The amino acid encoded by the codon is indicated. **c,** Correlation of the readthrough scores for the -1 codon mutants across the three genes. **d,** Mean readthrough difference between pairs of codons with Hamming distance of 1 (that is, single nucleotide difference) that encode for the same amino acid or pairs that encode a different amino acid, across genes. *P* value of the two-sided Wilcoxon signed-rank test between the same-AA and different-AA groups is shown. The top and bottom sides of the box are the lower and upper quartiles. The box covers the interquartile interval, where 50% of the data are found. The horizontal line that splits the box in two is the median. **e,** We used the *AQP4* and *MAPK10* double mutants to train models to predict readthrough based on the upstream sequence. The mean model coefficient size for each position upstream of the stop codon is shown. Coefficient units are in the underlying genetic trait scale, not as readthrough percentages (‘*Model design*’ Methods section). Bars show the standard error of the mean (SEM). **f,** Model coefficients colored by size. Missing mutations are colored in grey. **g,** Correlation of the model coefficients across genes, per position.

We first focused on the saturation mutagenesis of the -1 codon **(Fig. 4b)**, where mutants span a large dynamic range, likely reflecting the strong readthrough modulation exerted by the tRNA in the P-site. Consistent with this, several codons reduce readthrough to the levels of the second-stop-codon negative controls, including all four proline codons, all glycine codons except GGA, and the ACT (threonine) codon, amongst others.

We first compared readthrough across genes, and saw a high correlation across all three pairwise comparisons (*R* = 0.9, *R* = 0.8 and *R* = 0.72 across n = 61 codons, **Fig. 4c**), indicating that the -1 codon effect is largely independent of the surrounding sequence. A surprising exception is the arginine codon CGG, which is the 4th strongest *MAPK10* mutant, whereas in *OPRK1* and *AQP4* it ranks at positions 44 and 50, respectively, suggesting some degree of sequence context influence shaping the effect of this specific codon.

We then asked whether the effects of -1 codon mutations cluster by amino acid (AA) identity, and detected frequent differences between codons encoding the same AAs **(Fig. 4b)**. As two representative examples, in *MAPK10* the alanine and serine codons display 0.6%, 1.7%, 2.3%, 3.1% and 1%, 1.5%, 2%, 3.1%, 4.5%, 7.2% readthrough, respectively. Additionally, leucine, phenylalanine, glutamine and arginine also show large inter-codon differences. In contrast, other AAs show more similar profiles such as histidine, isoleucine, glycine and proline, where the last two are the AAs with the lowest readthrough across genes. The low readthrough of proline codons is potentially explained by proline having the slowest peptide bond formation rates among amino acids<u>(Pavlov et al. 2009; Johansson et</u> <u>al. 2011; Huter et al. 2017)</u>, which would reduce the likelihood of a readthrough event. To further explore the extent to which codon effects are driven by AA-dependent versus AA-independent mechanisms, we computed all readthrough differences across pairs of codons at 1 Hamming Distance (to control for higher nucleotide differences between different-AA vs same-AA codons) **(Fig. 4d)**. Overall, same-AA codons show more similar readthrough than different-AA codons (*P* = 7.38e-06, *P* = 1.41e-03 and *P* = 7.43-e06, for *AQP4*, *MAPK10* and *OPRK1*, respectively), suggesting that AA identity is partially driving the effects of the -1 codon. To identify additional determinants, we examined readthrough association with several tRNA-related metrics, including genomic tRNA copy number, tRNA abundance in the HEK293T cell line<u>(Behrens et al. 2021)</u> and the tRNA adaptation index (tAI)<u>(dos Reis et al. 2003; dos Reis et al. 2004)</u>, but no strong correlation was detected **(Supplementary Tables 2-4; Methods)**.

### Interpretable models for the upstream sequence

We next focused on the *AQP4 and MAPK10* double upstream mutants (we discarded the *OPRK1* triple mutants, see **Methods**) and fitted a simple additive model with sigmoidal global epistasis to obtain a coefficient for each position-codon pair (each observed an average of 3 times per gene), which allowed a direct comparison of the effects of each codon across the 6 upstream positions, using as an arbitrary reference the GCA codon **(Fig. 4f, Methods)**.

As expected, the -1 position shows the largest coefficients consistently across genes, supporting its stronger readthrough contribution compared to the other positions **(Fig. 4e,f)**. However, leaving the -1 codon unmodified and mutating the others can still yield strong readthrough changes **(Fig. 4a,f)**. For instance, among the 298 *MAPK10* variants carrying upstream mutations at positions other than the -1 codon, 32 exhibit readthrough levels below 1% (FDR = 0.05; one-sample T-test against a 1% threshold). Interestingly, the effect of mutations in -2 to -6 positions is substantially different across the two genes (mean *AQP4* vs *MAPK10* correlation across -2 to -6 position, *R* = 0.15), whereas in -1 it is highly similar (*R =* 0.85) **(Fig. 4g)**. We then asked whether codon effects are similar across positions within each gene by calculating all pairwise coefficients’ correlations across -1 to -6 positions for each gene separately, and observed a high discrepancy with mean *R* = 0.12 for both genes **(Extended Data Fig. 1f)**. Finally, we asked whether properties of the 6-residue peptide were predictive of readthrough. However, no strong correlations were observed for any of the AA nor tRNA-related properties, including after excluding the -1 codon (all abs(*R*) < 0.3, **Supplementary Tables 5-12; Methods**).

The pronounced gene specificity of the effects at positions -2 to -6, compared to the strong similarity observed at position -1, suggests that the upstream effects are mediated by mechanisms distinct from those operating at position -1 and that these mechanisms are more dependent on the surrounding sequence context.

### Chimeric variants cause strong readthrough

The strategies described above explored how mutations in the local sequence space surrounding each gene influence readthrough. However, we also tested how readthrough is affected by larger sequence perturbations by dividing each *WT* sequence into three upstream and three downstream blocks, each 5-9 nts long, and tested multiple combinatorial arrangements among the four genes (hereafter referred to as ‘chimeras’) **(Fig. 1c, 5a)**. As opposed to the previous sections, here we recovered most of the variants containing *OPRL1* blocks (only variants with the *OPRL1 WT* upstream sequence were lost, 12.5% of the total chimeric variants), and therefore included sequences from this fourth gene in downstream analyses. The length of the blocks depended on the *WT* sequence length, and was 5, 9, 5, 5 nts (upstream) and 5, 5, 9, 5 nts (downstream) for *AQP4*, *OPRK1*, *MAPK10 and OPRL1*, respectively. We evaluated all downstream block (DB) combinations in the context of the three *WT* upstream sequences, and conversely, all upstream block (UB) combinations in the context of the three *WT* downstream sequences plus the *OPRL1 WT* (n = 423 variants). This mutational strategy was designed to test for interactions between distal sequence elements. The readthrough distribution of the chimeric variants showed that a substantial fraction exceeded the readthrough levels of all *WT* sequences, reaching a maximum of 7.2% readthrough **(Fig. 1h)**.

### Strong epistatic interactions between sequence blocks

To dissect the contribution of the individual blocks, we first fitted first-order models, where each block has an independent additive effect on readthrough (again including a sigmoidal global epistasis function). These models yielded 10-fold cross-validated performance of R² = 0.52 for the downstream chimeras and R² = 0.88 for the upstream chimeras **(Fig. 5b)**. For the downstream blocks, the model coefficients showed that *AQP4* conferred the lowest readthrough in blocks 1 and 2 **(Fig. 5c)**. Instead in block 3, where effect sizes were substantially smaller overall, *AQP4* showed higher readthrough than *OPRK1* and *OPRL1*, although lower than *MAPK10*. In the upstream region, the effects were more concentrated in the block immediately preceding the stop codon. In this block, *OPRL1* produced the strongest effects, closely followed by *AQP4*, whereas *OPRK1* and *MAPK10* were associated with lower readthrough. Regarding the sequences in which all downstream or upstream blocks belong to the same gene (labelled as ‘*Upstream*’ and ‘*Downstream*’ in **Fig. 5c**), we observe two different patterns. Downstream sequences follow the same order as *WT*s, with readthrough of *AQP4* < *OPRK1* < *MAPK10* < *OPRL1*. Upstream, however, the order is reversed with *MAPK10* < *OPRK1* < *AQP4* (we did not recover variants with the OPRL1 upstream sequence).

**Figure 5.**
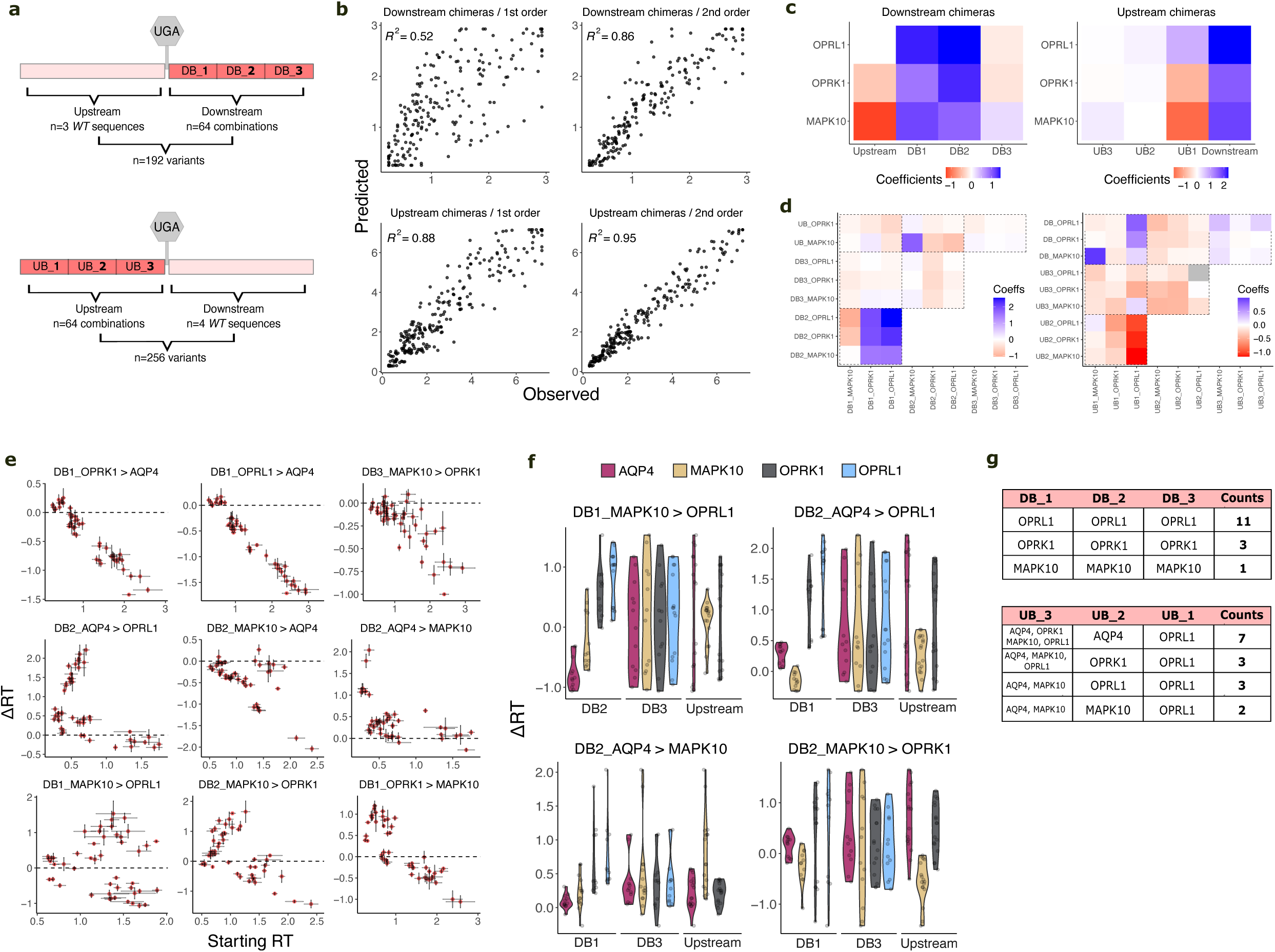
Chimeric variants can drive high readthrough. **a,** Mutagenesis strategy. We partitioned each sequence into three upstream and three downstream blocks (5-9 nts each, depending on the length of the *WT* sequence) and tested multiple combinatorial arrangements preserving the order (ie. block 1 always tested in position 1). All combinations of upstream blocks were evaluated in the context of the four *WT* downstream sequences, whereas all combinations of downstream blocks were tested with three *WT* upstream sequences (excluding *OPRL1 WT*). **b,** First-(*left*) and second-(*right*) order models for downstream (*top*) and upstream (*bottom*) mutants, cross-validated *R^2^* is shown. **c-d,** Coefficients of the first- (**c**) and second-(**d**) order models. Grey indicates the mutation was missing. Coefficient units are in the additive genetic trait scale, not in readthrough percentages. Blue and red indicate positive and negative effects relative to the reference sequence (*AQP4 WT*). **e,** Readthrough effects of the same block mutation across backgrounds, where the title indicates the mutation tested and each datapoint is a background. Axes indicate the readthrough of the background (Starting RT) and the readthrough change upon introducing the mutation (delta readthrough, ΔRT). Error bars indicate the standard deviation across the three replicates of the single mutant background (horizontal) and the double mutant (vertical). **f,** Similar to **e**, but with variants clustered according to their background sequence. Titles indicate the mutation tested, while subtitles denote the background blocks. Colours indicate the gene of each background block. Y axis shows the change in readthrough upon introduction of the mutation (ΔRT) in each background. **g,** Clustering the 15 chimeric variants with highest readthrough (RT > 5%) by their downstream (*top*) and upstream (*bottom*) blocks.

The first-order model provides good prediction of the upstream sequences (R² = 0.88) and fitting a second-order model that allows pairwise interactions (both between the upstream blocks and between the upstream blocks and the downstream *WT* sequences) slightly increased predictive performance (R² = 0.95) **(Fig. 5b)**. By contrast, downstream, the improved performance of a second-order model (R² = 0.52 vs 0.86 for 1^st^ and 2^nd^ order models, respectively) suggests the presence of strong epistatic interactions between blocks **(Fig. 5b)**. Indeed, analysis of the second-order coefficients reveals strong interactions, with positive and negative values indicating increases and decreases relative to the sum of individual effects **(Fig. 5d)**.

The strongest interactions were detected between the contiguous blocks DB1-DB2 and, to a lesser extent, between UB1-UB2 **(Fig. 5d)**. For instance, DB2_OPRL1 interacts negatively with DB1_MAPK10 but has a strong positive interaction with DB1_OPRL1. Positive interactions between blocks from the same gene (partial *WT* reconstitution) were also observed in other cases, such as between UB1_MAPK10 and DB_MAPK10. UB1_MAPK10 displays the lowest readthrough levels **(Fig. 5c)** and is rescued only by the *MAPK10* downstream sequence (DB_MAPK10) **(Fig. 5d)**. Similarly, DB2_MAPK10 shows strong specific epistasis with the upstream *MAPK10* sequence (UB_MAPK10). These two cases are particularly interesting because they represent upstream-downstream epistasis, showing how these two sequence regions interact to modulate readthrough. Moreover, the latter represents a long-range epistatic interaction, with 8-11 nts separating the two interacting blocks. Finally, we detected a strong pairwise interaction between upstream elements of *AQP4 and OPRL1*, where UB1_OPRL1 drives high readthrough only when combined with UB2_AQP4. In contrast, pairing it with UB2_OPRL1 (or UB2_OPRK1 or UB2_MAPK10) leads to a marked decrease in readthrough, indicating that the *OPRL1 WT* high-readthrough sequence **(Fig. 1b)** can be further optimized for readthrough.

To further explore interactions across blocks, we set out to analyse the differential effect sizes of mutations across sequence backgrounds by plotting the change in readthrough (ΔRT) for each mutation (block) across all backgrounds as a function of the background readthrough level (starting RT) **(Fig. 5e)**. We identified three different classes of blocks.

In class one, blocks show a strong relationship between ΔRT and the starting readthrough level, such that the baseline readthrough determines the magnitude of the effect and its direction remains constant, referred to as global epistasis **(Fig. 5e top)**. In the model, the sigmoidal global epistasis captures the readthrough lower bound (0%), causing ΔRT to scale with the starting readthrough.

In class two, blocks follow the same general trend but include a subset of outliers with magnitude epistasis, for which different ΔRT values are observed at the same initial readthrough level, but the effects on readthrough are always in the same direction **(Fig. 5e center)**.

Finally, the third class contains examples of sign epistasis **(Fig. 5e bottom)**. Here, both the direction and magnitude of ΔRT varies across backgrounds i.e. the same block introduced into different sequence contexts can both increase and decrease readthrough, depending on the background. For blocks in classes two and three, readthrough is shaped by both global and specific epistasis.

To combine the two previous analyses **(Fig. 5d,e)**, we grouped variants according to their background composition and compared their ΔRT distributions upon introduction of a given block **(Fig. 5f)**. For example, the effect of DB1_MAPK10>OPRL1 is strongly dependent on the identity of DB2. DB2 from *AQP4* results in negative ΔRT, DB2 from *OPRK1* and *OPRL1* in positive ΔRT, and DB2 from *MAPK10* exhibits a mixed effect. For DB2_AQP4>OPRL1, the identity of DB1 exerts a strong influence. The block has little to no effect in *AQP4* and *MAPK10*’s DB1, whereas it drives up to 2% ΔRT in the two other backgrounds. Finally, the DB2_AQP4>MAPK10 and DB2_MAPK10>OPRK1 mutations exemplify strong upstream-downstream interactions **(Fig. 5f)**. DB2_AQP4>MAPK10 produces a substantially larger ΔRT in the *MAPK10* upstream background than in the *AQP4* or *OPRK1* upstream backgrounds. Conversely, DB2_MAPK10>OPRK1 produces a negative ΔRT in the *MAPK10* upstream background but a positive ΔRT of up to 1.5% in the *AQP4* and *OPRK1* upstream backgrounds. Altogether, **Fig. 5c-e** shows the presence of strong epistatic interactions amongst different sequence elements.

Finally, we analysed the block composition of the chimeras with the highest readthrough (RT > 5%, n = 15 variants) and observed striking differences between upstream and downstream sequences **(Fig. 5g)**. Downstream, only block combinations that reconstitute the *WT* sequences are found, with 11, 3 and 1 occurrences for *OPRL1*, *MAPK10* and *OPRK1*, respectively. In contrast, the upstream sequences with highest readthrough are nearly all chimeric variants (13/15), with UB2_AQP4-UB1_OPRL1 present in 7/15 of these combinations, consistent with the second-order model in which these two blocks show a strong positive interaction **(Fig. 5d)**. Also in agreement with model coefficients, all 15 variants with the highest readthrough contain UB1_OPRL1.

## Discussion

Here we have reported a comprehensive mutagenesis of programmed stop codon readthrough in human genes, quantifying the effects of >1350 sequence variants in each of three endogenous readthrough contexts (*AQP4*, *MAPK10* and *OPRK1*), as well as 423 combinations of sequences from four different genes (chimeric variants). The dataset provides quantitative data and goes beyond prior studies restricted to testing tens of variants per gene, and allows us to systematically address questions about the sequence determinants, generalisability and genetic architecture of programmed readthrough.

In these three genes, the downstream CUAG motif is the key foundational sequence on which readthrough is built, with distinct sequence architectures layered around it further modulating readthrough. Downstream, readthrough modulating elements extend up to position +27, with particularly strong effects between positions +5 and +12. Upstream, the strongest effects are driven by the -1 codon, with milder modulation extending across several further codons. Both downstream and upstream individual mutations can abolish readthrough or double it relative to *WT,* and some -1 codon mutants increase it by up to ∼5-fold. Overall, this shows that distal sequence elements shape readthrough efficiency in human genes.

For single and double nucleotide mutant variants, the readthrough landscape is well captured by an additive model with sigmoidal global epistasis (R² = 0.84-0.95), with a small contribution from a small number of pairwise interactions, some spanning distances of up to 11 nts or bridging upstream and downstream regions.

However, an additional central finding of our study is the limited generalisability of mutational effects across genes outside of the core motifs (i.e. between much more diverged sequences than double mutants). The effects of mutations in the -1 codon and the CUAG motif generalise well, whereas mutations outside this immediate -3 to +4 window diverge across genes **(Fig. 2b**, **Fig. 3c)**. The -1 codon and the CUAG motif likely modulate readthrough through interactions with the ribosomal A and P sites and the eRF1/eRF3 complex, interactions that depend mostly on local sequence and are therefore largely independent of the surrounding context. By contrast, the gene-specific behaviour of more distal residues suggests a potential diversity of molecular mechanisms that may include secondary structure formation, diverse mRNA-ribosome contacts, recruitment of trans-acting factors, or changes in decoding kinetics.

Combining these results suggests an apparent paradox: within each gene, mutational effects combine additively, but this additivity breaks down when mutational effects are compared between very diverged sequences. This divergence of mutational effects with sequence divergence has been referred to as ‘epistatic drift’<u>(Park et al. 2022)</u> and reflects the accumulation of pairwise and potentially higher-order genetic interactions between mutations<u>(Escobedo et al. 2025; Beltran et al. 2025)</u>. The strong genetic interactions between sequence blocks from different genes in the chimeric constructs provide additional direct evidence for the importance of epistasis as sequences diverge. Mechanistically this epistatic drift likely reflects readthrough being promoted by distinct sequence motifs, secondary structures or other mechanisms in the different genes.

It appears therefore that the programmed readthrough sequence landscape is shaped by multiple local maxima: each programmed readthrough gene sits on (or near) its own fitness peak, defined by a particular combination of co-adapted upstream and downstream sequence features, separated from neighbouring peaks by valleys that require multiple interacting mutations to traverse.

More broadly, our approach establishes a quantitative framework for measuring and predicting programmed readthrough efficiency, and provides insights for future mechanistic dissection of the diverse molecular mechanisms that underlie this important form of information recoding.

---

**Extended Data Figure 1.**
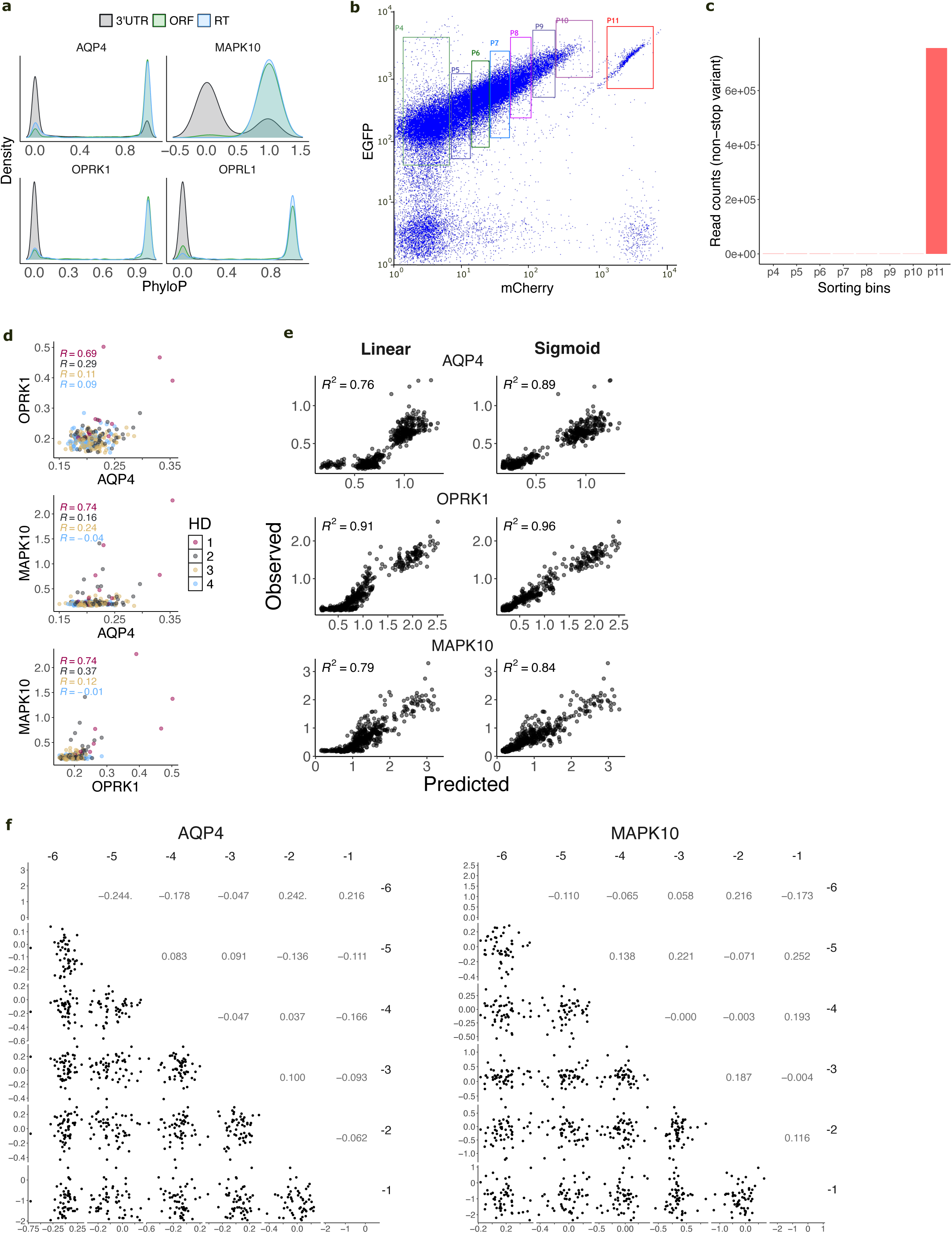
**a,** Distribution of PhastaCons scores across the open reading frame (ORF), readthrough extension (RT) and the rest of the 3’UTR across the *AQP4*, *MAPK10*, *OPRK1* and *OPRL1* genes. **b,** Fluorescence sorting strategy. EGFP+ cells were sorted in eight different bins according to mCherry fluorescence. **c,** Read counts distribution across sorting bins for the non-stop control variant. **d,** Correlation of readthrough scores for all CUAG-randomized variants across genes, clustered by mutation order (Hamming Distance (HD) to the CUAG WT sequence) Comparison **e,** Comparison of linear vs non-linear (sigmoid modelling global epistasis) models using the downstream sequence to predict readthrough. Cross-validated *R^2^* is shown. **f,** Correlations of the *AQP4* (left) and *MAPK10* (*right*) codon coefficients across the six positions upstream of the stop codon.

## Methods

### Ethics and consent

No specific ethics approval was required for this study.

### Readthrough reporter

Readthrough was measured using the dual fluorescent reporter plasmid previously described in Toledano et al. 2024<u>(Toledano et al. 2024)</u>. In brief, the plasmid drives expression of a single transcript carrying, in the 5’-3’ direction, the open reading frames (ORFs) of EGFP, two T2A sequences and mCherry. The library oligopool is inserted in frame between the two T2A elements. The T2As enable the fluorescent proteins to fold independently and shield them from potential effects of the variable sequence on folding and stability. Under standard translation, termination occurs at the stop codon of the library, preventing mCherry production. When readthrough occurs instead, elongation proceeds past this stop codon and continues until the mCherry stop codon, producing the mCherry protein along the way. Consequently, mCherry fluorescence scales with readthrough efficiency and serves as the assay readout. EGFP is used to exclude cells that either lack EGFP signal or show unexpectedly high levels of it. Such cells likely harbor aberrant cloning events, out-of-frame integration, promoter mutations, promoter silencing, or transcript-stability mutations, and could bias the assay if retained. The plasmid is compatible with genomic integration via the HEK293T landing pad system<u>(Matreyek et al. 2020)</u>, which guarantees single-variant integration per cell and thereby establishes a direct genotype-phenotype link. The vector carries BxBI-compatible attB sites that enable recombination into the landing pad of HEK293T cells. Once integrated into the landing pad locus, the ORF lies immediately downstream of a tetracycline-inducible cassette, so that expression is triggered upon doxycycline addition to the medium.

### Library cloning

Oligos were synthesised as an oligopool by Twist Biosciences, comprising the variable library region flanked by two constant sequences for PCR amplification. The pool was PCR-amplified for 14 cycles using primers oIT204 and oIT340 **(Supplementary Table 13)** and subsequently inserted between the EGFP-T2A and T2A-mCherry ORFs by Gibson Assembly. The assembled library was electroporated into Neb10 electrocompetent bacteria and propagated overnight in 50 mL of liquid culture. Library complexity and variant representation were estimated by plating an aliquot of the transformation reaction and extrapolating to the total transformant count. Individual colonies were Sanger-sequenced to verify the expected construct architecture and sequence diversity

### Stable cell line generation

Cell lines were generated in the HEK293T landing pad line developed by Matreyek et al. (TetBxB1BFP-iCasp-Blast Clone 12 HEK293T cells), which supports stable single-copy genomic integration of variants<u>(Matreyek et al. 2020)</u>. The library-containing reporter plasmid was co-transfected at a 1:1 ratio with the BxBI expression construct (pCAG-NLS-Bxb1, Addgene) into the HEK293T landing pad line in two T150 cm² flasks, using Lipofectamine 3000 according to the manufacturer’s instructions. This cell line carries a genomically integrated tetracycline-inducible cassette followed by a BxBI recombination site and a split rapalog-inducible dimerisable Casp-9. Cells were cultured in DMEM containing 10% tetracycline-free FBS, without antibiotics. Two days post-transfection, doxycycline (2 μg/mL, Sigma-Aldrich) was added to drive expression of either the integrated library (in recombined cells) or the iCasp-9 protein (in non-recombined cells). Twenty-four hours later, rimiducid (AP1903, Selleckchem) was added at 10 nM. Successful recombination shifts iCasp-9 out of frame, abolishing its expression. Non-recombined cells, in contrast, continue to express iCasp-9, which dimerises in the presence of rimiducid and induces apoptosis. Twenty-four hours after rimiducid treatment, the medium was replaced with DMEM supplemented with doxycycline, and cells were expanded over the following days to generate sufficient material for downstream experiments and cryopreservation.

### Fluorescence activated cell sorting (FACS)

Cells were cultured on standard tissue-culture plates in DMEM supplemented with 10% tetracycline-free FBS and no antibiotics. They were passaged before reaching confluency to preserve cell health. For harvesting, cells were detached with trypsin, pelleted by centrifugation, and washed with PBS. For the sort-seq experiments, transcript expression was induced by treating cells with 2 μg/mL doxycycline for 48 h prior to sorting. Large numbers of cells were used to ensure that each variant was represented more than 100 times in the population.

Sorting was carried out on a BD Influx™ Cell Sorter, with analysis in BD FACS™ Software (1.0.0.650). Gating was applied sequentially: forward and side scatter area to retain intact cells, forward scatter width and height to exclude aggregates, and DAPI staining to keep only live, recombined cells. EGFP and mCherry were excited with 488 nm and 561 nm lasers and recorded through 530/40 BP and 593/40 BP filters, respectively. EGFP-positive cells were then sorted by mCherry signal into eight populations **(Extended Data Fig. 1b)**. Readthrough scores for stop codon variants were calculated from read counts across P4-P10, whereas P11 was reserved for determining the distribution of the non-stop control variant. As anticipated, the control variant was more than 1000-fold enriched in P11 relative to the other gates **(Extended Data Fig. 1c)**. 400,000 cells were collected from each population, except for the smaller P10 and P11 fractions, where 250,000 and 150,000 cells were sorted, respectively; yielding a total of 2.8 million sorted cells per replicate (∼460 cells per variant). The proportion of cells in each bin was used as a normalisation factor in the sequencing analysis (see *Sequencing data processing* section below). All experiments were performed in biological triplicates.

### DNA extraction

Sorted cells were centrifuged at 1200rpms for 3mins, and the pellet was used to extract genomic DNA following the DNeasy Blood&Tissue Kit (Qiagen), and resuspended in 75 ul of Milli-Q water.

### Sequencing library preparation

The sequencing libraries were prepared through three sequential PCR reactions. The aim of the first PCR was to selectively amplify the library fragment from the genomic DNA pool while avoiding amplification of any residual plasmid carried over from transfection. To achieve this, a forward primer (oIT314) annealing within the landing pad, outside the recombined region, was paired with a reverse primer (oIT205) annealing at the 3’ end of the library fragment. Plasmid DNA lacks the forward-primer annealing site and is therefore not amplified. The second PCR (PCR2) introduces part of the Illumina adapter sequences and increases the nucleotide complexity of the first sequenced bases by adding frame-shift bases between the adapters and the region of interest. The third PCR (PCR3) appends the remaining Illumina adapter sequence and the demultiplexing indexes. All PCRs were performed with Q5 Hot Start High-Fidelity DNA Polymerase (New England Biolabs) following the manufacturer’s protocol.

For PCR1, the entire genomic DNA recovered from each bin was used as template, together with 25 pmol of primers oIT205 and oIT314. Reactions were run with a 66 °C annealing temperature, a 1-minute extension, and 25 cycles. Because large amounts of genomic DNA inhibit the PCR, each sample was split across 8 reactions arrayed in 96-well plates. The 8 reactions per sample were subsequently pooled and purified with the MinElute PCR Purification Kit (QIAGEN) according to the manufacturer’s protocol, and the DNA was eluted in 20 μL of Milli-Q water.

PCR2 was performed using 2 μL of the PCR1 product as template, together with 25 pmol of the pooled frame-shift primer sets oIT_ILL_204_mix and oIT_ILL_205_mix **(Supplementary Table 13)**. Cycling conditions were 66 °C annealing, 15 seconds of extension, and 8 cycles in total. Reactions were then purified per sample with the MinElute PCR Purification Kit (QIAGEN) following the manufacturer’s protocol, and the DNA was eluted in 10 μL of Milli-Q water.

For PCR3, 2 μL of the PCR2 product served as template, and the remaining portions of the Illumina adapters were appended to the library amplicon. Each sample was amplified with a distinct combination of barcoded primers (oIT_GJJ_1J and oIT_GJJ_2J; **Supplementary Table 13**), enabling pooling and downstream demultiplexing after sequencing. PCR3 was run for 8 cycles with a 62 °C annealing temperature and a 25-second extension, after which an aliquot was resolved on a 2% agarose gel for quantification. Samples were then pooled in equimolar amounts, run on a gel, and purified with the QIAEX II Gel Extraction Kit. The purified amplicon pool was sequenced on an Illumina NextSeq2000 (P2 flowcell, 150 bp paired-end reads) at the CRG Genomics Core Facility.

### Sequencing data processing

FastQ files from paired-end sequencing of all experiments were processed with DiMSum (v.1.3)<u>(Faure et al. 2020)</u> (reference https://github.com/lehner-lab/DiMSum) to obtain the read counts for each variant. DimSum applies stringent quality filters to discard low quality reads, reads with sequencing errors, etc., to ensure that only high-quality reads are used for downstream analysis.

The DimSum output read count table is used to estimate the distribution of each variant among the different sorting gates. Since gates represented different percentages of the general population, they had to be sorted for different times in order to get similar number of cells in each bin, forcing us to calculate the mCherry distributions in relative numbers as follows:

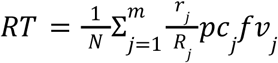

The distribution of each variant is estimated by calculating its proportion of reads in each sorting gate (*j*). All reads of a given variant in a given sorting gate (*r_j_*) are a) divided by the total number of reads of that gate (*Rj*) yielding a normalized reads value b) assigned a first fixed value (*pc*) corresponding to the % of cells of the total population that belong to that gate and c) a second fixed value (*fv*) corresponding to the mean mCherry signal of the gate relative to the mCherry value of the P11 population. Finally, in step d), the score averaged by the total number of normalized reads across gates (N). In steps a) and b) we are simply calculating the percentage of reads of the total population belonging to each variant in each gate 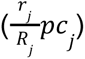. In step c) and d), we are using these scores to calculate the mean mCherry value of each variant, where mCherry is already in a percentage scale (% of the non-stop P11 population). This yields a readthrough percentage for each variant, which represents the percentage of expression of that variant relative to a non-stop control variant. Standard deviation from the three replicates is used as the error measure. Readthrough scores for all variants are shown in **Supplementary Table 1**. Experiments were performed in biological triplicates and 5036 high-confidence variants (≥100 reads) were recovered in each experiment. We used an unusually high coverage threshold (≥100 reads) because the sample was oversequenced (mean of 5,000 reads per variant).

### Model design

We fitted models analogous to those previously used to characterise drug-induced readthrough<u>(Toledano et al. 2024)</u>, in which mutations combine additively to determine an underlying genetic trait (the ‘readthrough potential’), which is then mapped onto the observed readthrough through a sigmoidal function. We used the same model formulation for all models described throughout the paper (downstream sequence, upstream sequence and chimeras). As an example, the formula used for the *AQP4* downstream sequence model is shown below:

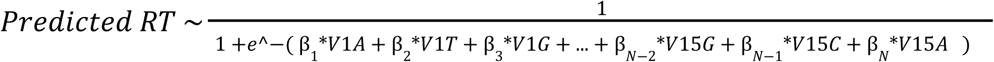

To assess the contribution of the sigmoidal non-linearity we also fitted purely linear models, which consistently performed worse and produced residuals with larger systematic bias **(Extended Data Fig. 1e)**. Models were trained on 90% of the dataset and tested on the remaining 10% of variants (held-out set). This procedure was repeated 10 times, and the mean R² across cross-validation rounds was used as the performance metric.

For the upstream mutants, we trained models only for double mutants (*AQP4* and *MAPK10*) but not for triple mutants (*OPRK1*). The higher mutation order of the triple mutant library means each variant carries more co-occurring mutations, making the contribution of any individual mutation harder to disentangle from those of its partners and yielding unreliable per-codon coefficient estimates (at this library size).

### Individual measurement of *WT* variants

To ensure that *AQP4*, *OPRK1*, *MAPK10* and *OPRL1* sequences triggered readthrough in our system, we individually measured them by generating stable HEK293T cell lines expressing a single copy of each sequence from the landing pad. We used the same settings as for the library, where cells were treated with 2ug/mL doxy 48h before fluorescence quantification. EGFP and mCherry signal was quantified using 530/40 BP and 593/40 BP channels in BD LSRFortessa™ Cell Analyzer. EGFP+ cells were used to calculate the readthrough efficiency, multiplying the mean mCherry intensity by the percentage of mCherry+ cells, and finally normalizing by the expression of the non-stop variant:

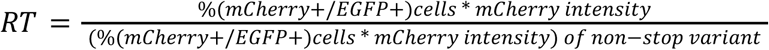

### tRNA Adaptation Index (tAI)

The tRNA Adaptation Index (tAI) is a measure of translational efficiency which takes into account the intracellular concentration of tRNA molecules and the efficiencies of each codon-anticodon pairing<u>(dos Reis et al. 2003; dos Reis et al. 2004)</u>. The pairing affinity of each codon-anticodon is specific to each species. The human specific tAI indexes were downloaded from the STADIUM database as of January 2023<u>(Yoon et al.</u> <u>2018)</u>. tAI for a given sequence was calculated as the mean tAI across all codons of the sequence.

### tRNA abundance and copy number

Both metrics were retrieved from Behrens et al.<u>(Behrens et al. 2021)</u>. tRNA abundance data was generated in HEK293T cells.

### Aminoacid features

We correlated readthrough scores with a set of 548 amino acid property scales sourced from the AAindex database (https://www.genome.jp/aaindex/) <u>(Kawashima et al. 2008)</u>, and listed them in **Supplementary Tables 5 and 7**.

### Statistics & Reproducibility

Statistical tests were performed in R (v4.4.2) using RStudio (2024.12.0+467). Statistical significance between pairs of variants was assessed using T-tests across three biological replicates. *P* values were adjusted for multiple testing using FDR correction. Variants with <100 reads were not used for analyses.

## Acknowledgements

This work was funded by Wellcome (Grant reference: 220540/Z/20/A, ‘Wellcome Sanger Institute Quinquennial Review 2021-2026’), European Research Council (ERC) Advanced Grant 883742 and Proof of Concept Grant 101290597, the Spanish Ministry of Science and Innovation (BFU2017-89488-P, EMBL Partnership, Severo Ochoa Center of Excellence), Agencia de Gestió d’Ajuts Universitaris i de Recerca (AGAUR, 2021 SGR 01226) and the CERCA Program/Generalitat de Catalunya.

## Author contributions

I.T., F.S. and B.L. conceived the project and designed the experiments. I.T. performed the experiments and analyzed the data. I.T. and B.L. wrote the manuscript.

## Data availability

All DNA sequencing data have been deposited in the Sequence Read Archive with accession <u>PRJNA1494055</u>. All readthrough measurements are provided in Supplementary Table 1.

## Code availability

All scripts used in this study are available at: https://github.com/lehner-lab/Programmed_readthrough

## Competing Interests

B.L. is a founder and shareholder of ALLOX and on the Scientific Advisory Board of Metaphore Biotechnologies. The remaining authors declare no competing interests.

